# Flatfish intestinal microbiota depend on various host traits, and vary with sediment type and bottom trawling effort

**DOI:** 10.1101/2025.05.30.657041

**Authors:** Michelle Gwinner, Holger Haslob, Hermann Neumann, Sahar Khodami, Peter J. Schupp, Guido Bonthond

## Abstract

The intestinal microbiota of fishes support digestion, nutrient uptake and play an important role in the immune system, development and reproduction. Flatfish live in close contact with the seafloor, and are particularly exposed to anthropogenic disturbances such as bottom trawling. Bottom trawling impacts the ecosystem in various ways and it recent evidence indicates that the microbial composition and diversity in marine sediments varies with fishing intensity. It is presently unknown whether this trawling signal applies to the seafloor alone, or may also extend to microbiota of marine holobionts inhabiting it, such as flatfishes. Here, three flatfish species (*Buglossidium luteum*, *Limanda limanda* and *Pleuronectes platessa*) were sampled across the southeastern North Sea. We characterized the intestinal microbiota using 16S rDNA metabarcoding of 162 individuals, and disentangled how intestinal microbial composition and diversity are jointly shaped by various host traits (species, sex, age, weight, and condition factor) and environmental factors (sediment type and trawling intensity). Intestinal diversity varied among species and changed with age, weight and sediment type. Community composition was dependent on species, age, condition factor and sediment type. In addition, we found that trawling intensity explained shifts in intestinal microbial community composition, suggesting that the known impacts of bottom trawling on the benthic environment may cascade to intestinal microbiota of flatfish. Our findings provide important insight into host-microbiota interactions in marine ecosystems and highlight the interplay between host traits and environment as drivers of intestinal microbial diversity and community composition in flatfish.

**Highlights:**

- Flatfish microbiota are shaped by host traits and environmental factors
- Host species and age affect intestinal microbial diversity and composition
- Intestinal microbiota vary along a bottom trawling intensity gradient
- Bottom trawling impacts may cascade from sediments to fish intestinal microbiota

## 1. Introduction

Comprising a diverse and complex ecosystem of bacteria, archaea, fungi, protozoa, and viruses, the intestinal microbiota play a critical role in animal physiology in general, and in fish particularly (Arun & Midhun 2023; Rombout *et al*. 2011). For instance, the intestinal microbiota influence immune functions, homeostasis, food digestion, growth and reproduction (Nayak 2010; Nie *et al*. 2017; Ray *et al*. 2012; Rolig *et al*. 2017; Tarnecki *et al*. 2017; Wu *et al*. 2012). In humans, where intestinal microbiota have been studied most extensively, healthy intestinal microbiota have even been shown to enhance brain health and stress response (Dinan & Cryan 2016). Due to their shorter generation times and higher genetic variability, microbiota can respond more rapidly to environmental changes, thereby influencing the adaptive responses of their marine hosts (Leray *et al*. 2021; Wilkins *et al*. 2019).

The factors that shape the composition and diversity of these microbial communities through space and time are less well understood, but appear to be habitat, species and tissue specific (Ghotbi *et al*. 2022; Huang *et al*. 2020; Kanika *et al*. 2025; Li *et al*. 2017), and correlate with variables such as diet, age, pollutants, salinity or temperature (Adamovsky *et al*. 2018; Bolnick *et al*. 2014; Lozupone & Knight 2007; Nayak 2010; Ringoe & Birkbeck 1999).

As the largest and most diverse group of vertebrates comprising more than 30,000 species (Fan *et al*. 2020), fish play an essential role in a variety of aquatic ecosystems. Notably, they contribute to maintaining the biodiversity, stability and resilience of those ecosystems. Further, as a fundamental part of aquatic food webs, they ensure the recycling and transport of diverse nutrients, even between aquatic and terrestrial ecosystems (Holmlund & Hammer 1999). As consumers in marine food webs, fish contribute to the biological carbon pump by transporting carbon to the seafloor, thereby helping to mitigate the impacts of climate change (Saba *et al*. 2021). Beyond providing fundamental ecosystem services, fish account for important economic value and contribute to global nutrition (FAO 2024; Viana *et al*. 2023). Despite the high species diversity, only a small fraction is used commercially as food fish (FAO 2022). For instance, in the North Sea, home to approximately 197 fish species, only a small number of species accounts for the majority of total landings (Froese & Pauly 2024; ICES 2022). Demersal fish, such as flatfish, account for a significant proportion of these landings, highlighting their importance in the region’s fisheries (ICES 2022; Link *et al*. 2002). With the rapid decline of fish populations and the collapse of entire commercially exploited stocks, such as the North Sea herring (*Clupea harengus*) in the 1970ies, gaining a deeper understanding of population dynamics and fish health is increasingly important (Dickey-Collas *et al*. 2010; FAO 2024; Zhu *et al*. 2021).

Fishing practices are known to impact the marine environment, with bottom trawling being the most pervasive anthropogenic disturbance of seabed habitats (ICES 2022; Kaiser *et al*. 2002). Shallow shelf sea regions are especially subject to trawling, with the central and south-eastern North Sea being among the most heavily trawled areas in the world (Amoroso *et al*. 2018). Besides the removal of flatfish directly, trawling may also indirectly impact the remaining flatfish populations, by altering the communities of seabed organisms that partially form their diet, or by resuspending the upper sediment layers, stirring up fine particulate matter and contaminants (Kaiser *et al*. 2002). Furthermore, recent evidence suggests that trawling may also alter the microbiota of the sediments (Bruce *et al*. 2022; Bonthond *et al*. 2023). Both diet and sediment serve as crucial microbial sources for the microbial communities associated with flatfishes and other seafloor-inhabiting holobionts, playing a primary role in shaping the composition and diversity of their intestinal microbiota. However, whether trawling impacts on flatfish diet and sediment microbiota may be indirectly affecting the microbiota of seafloor dwelling holobionts, including flatfishes, remains unknown.

To gain better insight into the flatfish intestinal microbiota and improve our understanding of wild flatfish populations, we analyzed the intestinal microbiota of three flatfish species (*Buglossidium luteum*, Risso 1810; *Limanda limanda,* Linnaeus 1758; and *Pleuronectes platessa*, Linnaeus 1758) that were collected in the southeastern North Sea. Using 16S rDNA metabarcoding, we characterized the prokaryotic communities of 162 fish. Subsequently, we used uni- and multivariate generalized linear models to test the predictive potential of different variables on the structure and diversity of the intestinal microbiota, each representing individual hypotheses. This included a combination of host variables (i.e., species, sex, condition factor, age and weight), and environmental variables (i.e., sediment type and trawling intensity). In addition, we analyzed differential abundances associated with these variables, to identify microbial markers that potentially fulfil important roles in the flatfish intestinal microbiota.

## 2. Material and Methods

### 2.1 Sampling

Flatfishes were sampled during an expedition with the German Research Vessel Solea from 07.05.2021 to 18.05.2021 (Cruise number: SOL791). The expedition took place in the Natura 2000 area in the Southeastern North Sea (Figure 1) and was carried out within the frame of a systematic survey on demersal fish and epibenthic fauna in the southeastern North Sea. A 2-meter beam trawl with a 20 mm mesh size outer net and 4 mm mesh size codend, and a 7-meter beam trawl with a 70 mm mesh size forenet and a 20 mm mesh size codend were used.. At each station, the two-meter beam trawl had a towing time of five minutes on ground, while the seven-meter beam trawl remained on the bottom for 15 minutes. Flatfish were caught exclusively during the day and for the present study the species *Pleuronectes*□*platessa* (European plaice), *Limanda*□*limanda* (Common dab) and *Buglossidium luteum* (Solenette) were selected from the catch. When possible, one male and one female were randomly selected at each station. The fish were placed in plastic bags and frozen at 20□°C on board. After the cruise, the fish were transferred to the laboratory in cooling boxes, where they were again stored at 20□°C. In total 183 individuals of the species *P.*□*platessa*, *L.*□*limanda* and *B.*□*luteum* were analyzed. Prior to dissection of fish individuals, the sex of the thawed fish was re-determined. Then, the length of the fish was measured from the head to the end of the tail fin and the weight was recorded. Scissors, forceps, and scalpels were cleaned and sterilized with a burner before the dissection of each fish and all procedures were carried out in a sterile hood.

**Figure 1:**
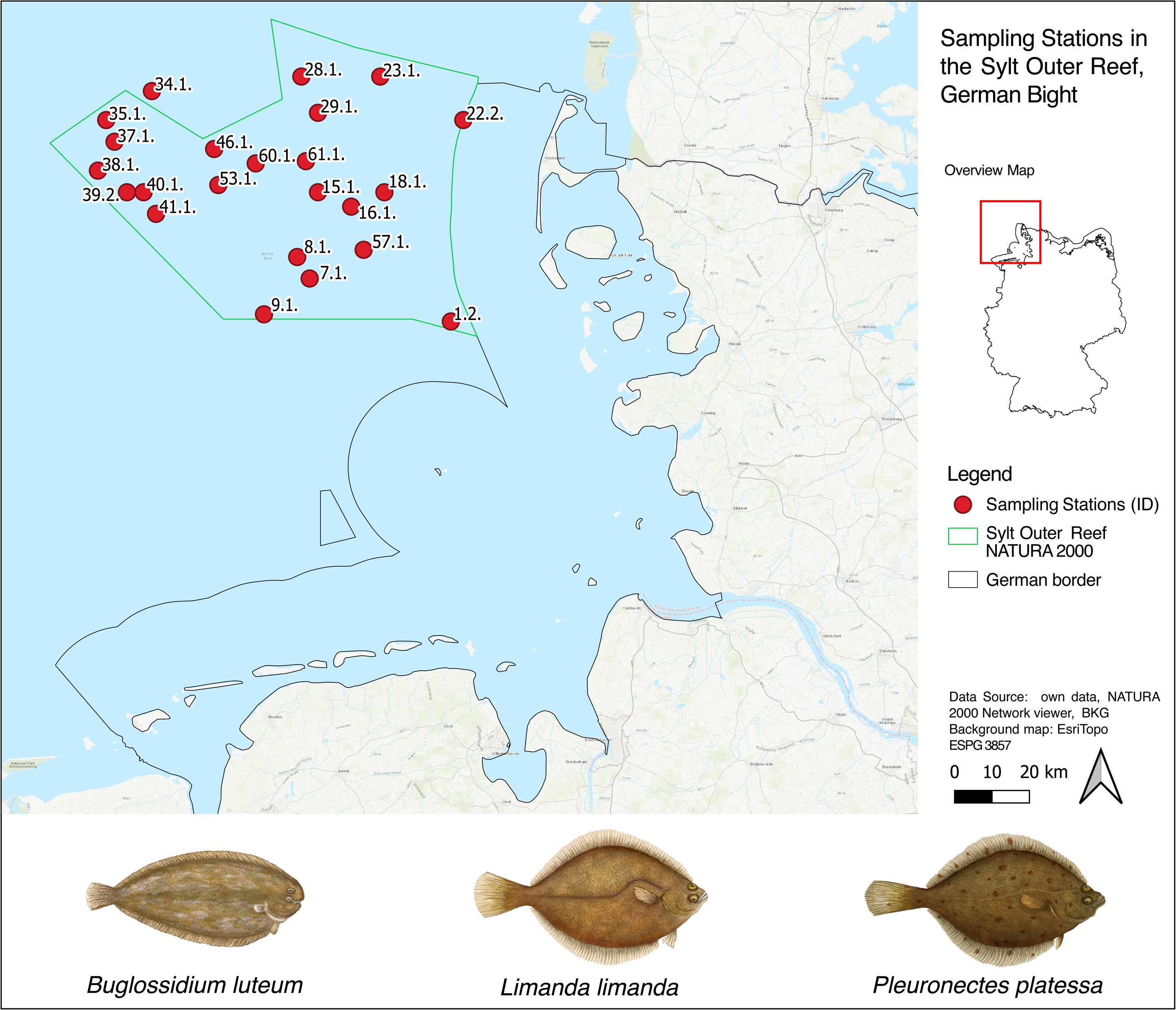
Sampling stations in the Sylter Outer Reef area (green outline) in the Southeastern North Sea and target species below. Red circles highlight the sampling stations where the fish were caught. Drawings of the species studied here can be seen below the map.

To determine the age of the fish, the otoliths were removed, cleaned and stored in individually in tubes, and the age was determined at the Thünen Institute of Sea Fisheries (Bremerhaven, Germany). Subsequently, the stomach, intestines, liver and in females also the gonads were removed, and their weight was measured individually. Intestines were then collected in separate tubes and stored at -20□°C for DNA extraction.

### 2.2 DNA extraction and library preparation

To allow optimal DNA extraction from intestinal prokaryotes, intestines were cut into small fragments with sterile scissors, resulting in a homogeneous suspension. From this suspension, subsamples of 100 to 125 mg were used for DNA extraction with the ZYMO Quick-DNA™ Fecal/Soil MicroPrep Kit (D6012; ZYMO Research) according to the manufacturer’s protocol. To verify potential contamination during the extraction process, DNA was also extracted from several blanks. Amplicon library preparation followed the two-step PCR protocol from (Gohl *et al*. 2016). The 16S-V4 region was amplified using the primers 515F (S-*-Univ-0515-a-S-19) and 806R (S-D-Arch-0786-a-A-20, Klindworth et al., 2013), using the Phusion Green Hot Start II High-Fidelity PCR Master Mix (ThermoFisher Scientific), and 1□µl of DNA template per reaction. PCRs were also conducted on Mock communities (ZYMO-D311) and negative controls. The program of the first PCR included an initial denaturation step of 98□°C for 3:00 min, followed by 30 cycles of 98□°C for 0:30 min, 55□°C for 0:30 min and 72□°C for 0:30 min and a final elongation step of 72□°C for 5:00 min. PCR products were run through a 1% agarose gel electrophoresis and samples without a visible product underwent 5 additional PCR cycles. Then, PCR products were diluted 1:100 and used as templates for the second PCR, which was conducted with the same reagents and the indexing primers from Gohl et al. (2016). This PCR included 10□cycles but was otherwise identical to the PCR□1. The final amplicons were then normalized, purified and pooled using sequelPrep plates (ThermoFisher Scientific). The sequencing, on the Illumina MiSeq platform with the MiSeq Reagent□Kit□v3 (600-cycle), was done in the German Centre for Marine Biodiversity Research (DZMB; Wilhelmshaven, Germany).

### 2.3 Data preparation and processing

The raw sequencing reads were quality filtered and clustered into OTUs based on the 97% similarity criterion, using the OptiClust algorithm (Westcott & Schloss 2017) using the software MOTHUR (v.1.48.0, Schloss et al., 2009) and classified with the SILVA database (v138.1, Quast et al., 2012). Finally, samples were purged of reads classified as chloroplasts, mitochondria or eukaryotes, as well as those without a domain level classification, and excluded from downstream analyses if the remaining total read count was below 1000. The final dataset was rarefied by averaging 100 replicated count tables that were subsampled to 3500 read counts.

### 2.4 Statistical analysis

All analyses were performed in R (v4.4.2, see for rendered scripts https://github.com/gbonthond/flatfish_microbiota). We considered a total of eight host and environmental variables. The host variables included species, sex, age (in years), length (in cm), total weight (in grams) and Fulton’s condition factor (total weight/length³, Nash et al., 2006). The weight and age were transformed with a natural log and the median grain size with a log-base 2. The environmental variables that were considered for the analysis were the median grain size (in µm) as a proxy for habitat type and the trawling intensity in swept area ratio (SAR) per year. Median grain size data was obtained from the NOAH Habitat atlas portal (NOAH 2015). As a measure for the trawling intensity, we used fishing intensity of the subsurface (≥2 cm) penetrating gears from the OSPAR data & information management system (OSPAR 2017). Based on spearman correlation coefficients, which revealed a strong correlation between length and weight (Spearman’s rank coefficient = 0.99, Figure S1, and see Figures S2-5 for more details on the host variables), it was decided to use the variable weight as an indicator of size and exclude length from downstream analyses.

To evaluate the impact of the remaining predictors on the diversity of the intestinal microbial community, we calculated the OTU richness and the effective number of OTUs (Jost 2006). On both responses, we fitted linear mixed models (LMMs) using the R package lme4 (v1.1-35.5, Bates et al., 2015). First a global model was fitted with the same structure as used for the PERMANOVA (i.e., the main effects and all possible interactions with the factor species identity) and with the station identity as a random effect. Model assumptions were assessed using diagnostic plots. Model selection was then performed for both response variables by comparing the global model to all possible simpler models, keeping the random effect fixed. The best model was selected based on the corrected Akaike Information Criterion (AICc).

The community composition of the intestinal microbiota was analyzed with PERMANOVA (Anderson 2001) with the adonis2() function in the R package vegan (v2.6.8 (Oksanen *et al*. 2013), based on Bray Curtis distances and 9999 permutations. To test whether predictors had general effects applicable to all species, as well as species-specific effects, all seven predictors were included along with all possible second order interactions with the factor species identity. To visualize community similarity patterns, we conducted non-metric multidimensional scaling (nMDS) based on Bray Curtis distances, using the R package vegan (v2.6.8).

Subsequently, a differential abundances analysis was conducted to identify microbial markers for each of the predictors. First, the OTU dataset was reduced to OTUs with at least 1% occurrence and at least 0.1% relative abundance. Second, a multivariate generalized linear model (mGLM) was fitted on the reduced community matrix with the package mvabund (v4.2.1, Wang et al., 2012). This model included all seven predictors and assumed a negative binomial distribution. Third, the estimated model coefficients were used to identify marker OTUs for each predictor, with coefficients considered significant if the respective 95% confidence intervals did not overlap with zero. To identify species-specific OTUs, the mGLM was fitted three times, each time using a different species as the reference level, allowing us to obtain pairwise estimates of species differences. The two coefficients and standard errors of the differences with both other species were then pooled to obtain estimates and confidence intervals for species-specific OTUs.

## 3. Results

After quality filtering a total of 162 samples remained. These belonged to 65 individuals of the species *B.*□*luteum*, 51 of *L.*□*limanda*, and 46 of *P.*□*platessa*, and included 96 females and 66 males. The rarefied community matrix counted 12,676 OTUs. Pseudomonadota is the most abundant phylum followed by Actinomycetota, Planctomycetota, Bacillota, and Verrucomicrobiota. At the family level, Pirellulaceae, Ilumatobacteraceae, Paracoccaceae, Hyphomicrobiaceae, and Actinomarinales_uncultured were the five most abundant groups, accounting for 36.7% of the total OTU abundance (Figure 2).

**Figure 2:**
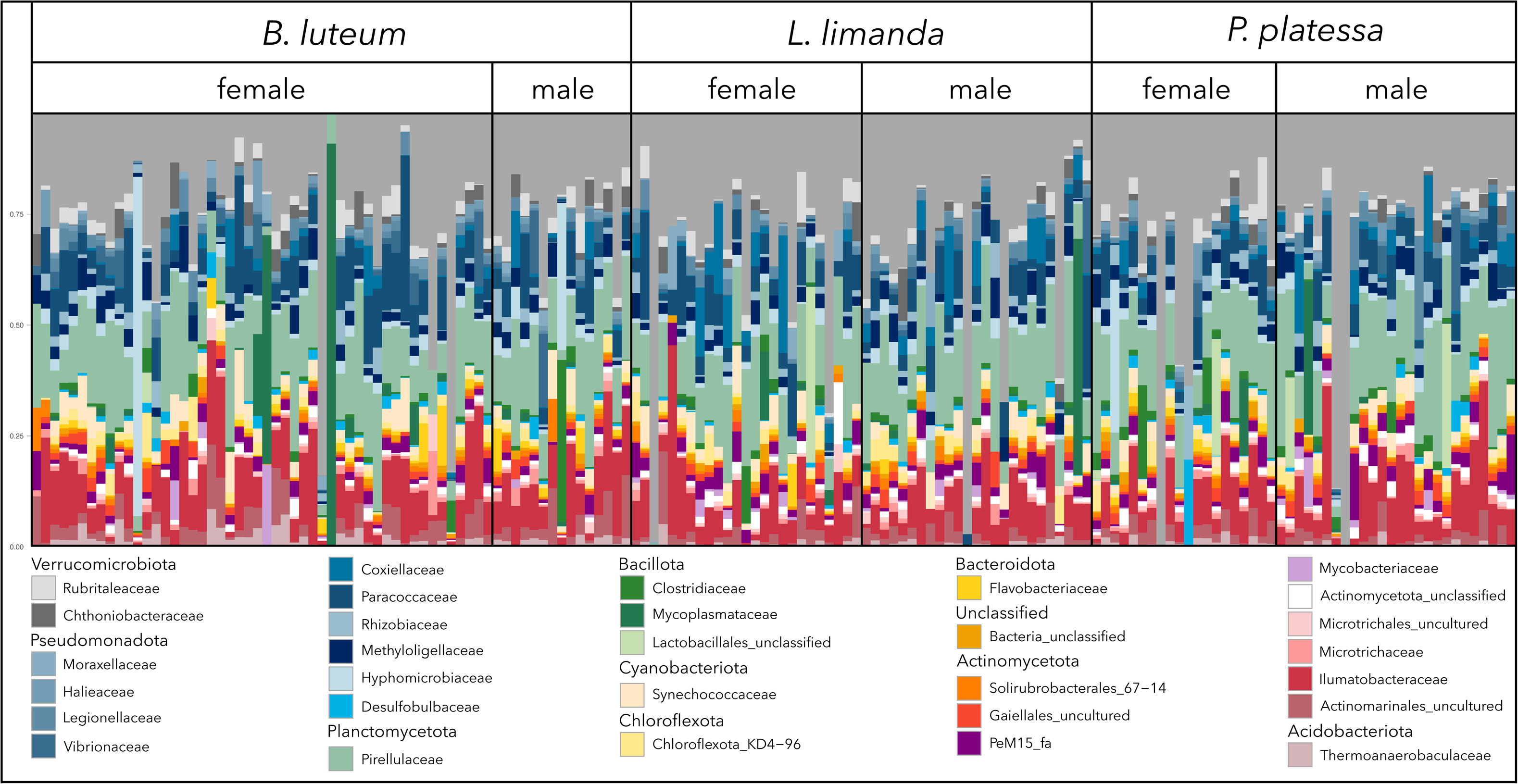
Stacked bar plot of the 30 most abundant prokaryotic families across all samples, sorted by species and sex. Each color represents a family, which are sorted by phylum. The length of the bars corresponds to the proportional abundance in the microbiome of a fish’s intestine.

While the intestinal microbiota were generally dominated by a combination of taxa from the phyla Pseudomonadota, Planctomycetota and Actinomycetota, some samples deviated from this pattern and only counted few taxa (e.g., Mycoplasmataceae or Hyphomicrobiaceae), or taxa that were not among the 30 most abundant families.

### 3.1 Diversity

From the comparison of models fitted on the effective number of OTUs (ENO) the best model yielded only trawling intensity as informative predictor (Table S1). However, the effect of trawling intensity on ENO was not significant (χ²_1_ = 3.06, *p* = 0.080, Table S2) and the model explained < 2% of the variation (R^2^m = R^2^c = 0.019). In contrast, for OTU richness, the best model (R^2^m = 0.151, R^2^c = 0.225) included the variables species (χ²_2_ = 7.33, *p* = 0.026), age (χ²_1_ = 5.05, *p* = 0.025), weight (χ²_1_ = 3.89, *p* = 0.049), the median grain size (χ²_1_ = 7.77, *p* = 0.005) and the interaction between the median grain size and species (χ²_2_ = 6.74, *p* = 0.034, (Table S1-S2). Post-hoc pairwise comparisons (adjusting *p*-values to control for familywise error rates with the Holm method) among species, revealed significant differences in OTU richness between *B.*□*luteum* and *P.*□*platessa* (t_167_ = -2.525, *p* = 0.030) and between *L.*□*limanda* and *P.*□*platessa* (t_158_ = -2.605, *p* = 0.030), and identified that of species-specific changes in OTU richness with the median grain size, only *L.*□*limanda* was significant (t_116.9_ = -3.209, *p* = 0.005, Figure 3, Table S3).

**Figure 3:**
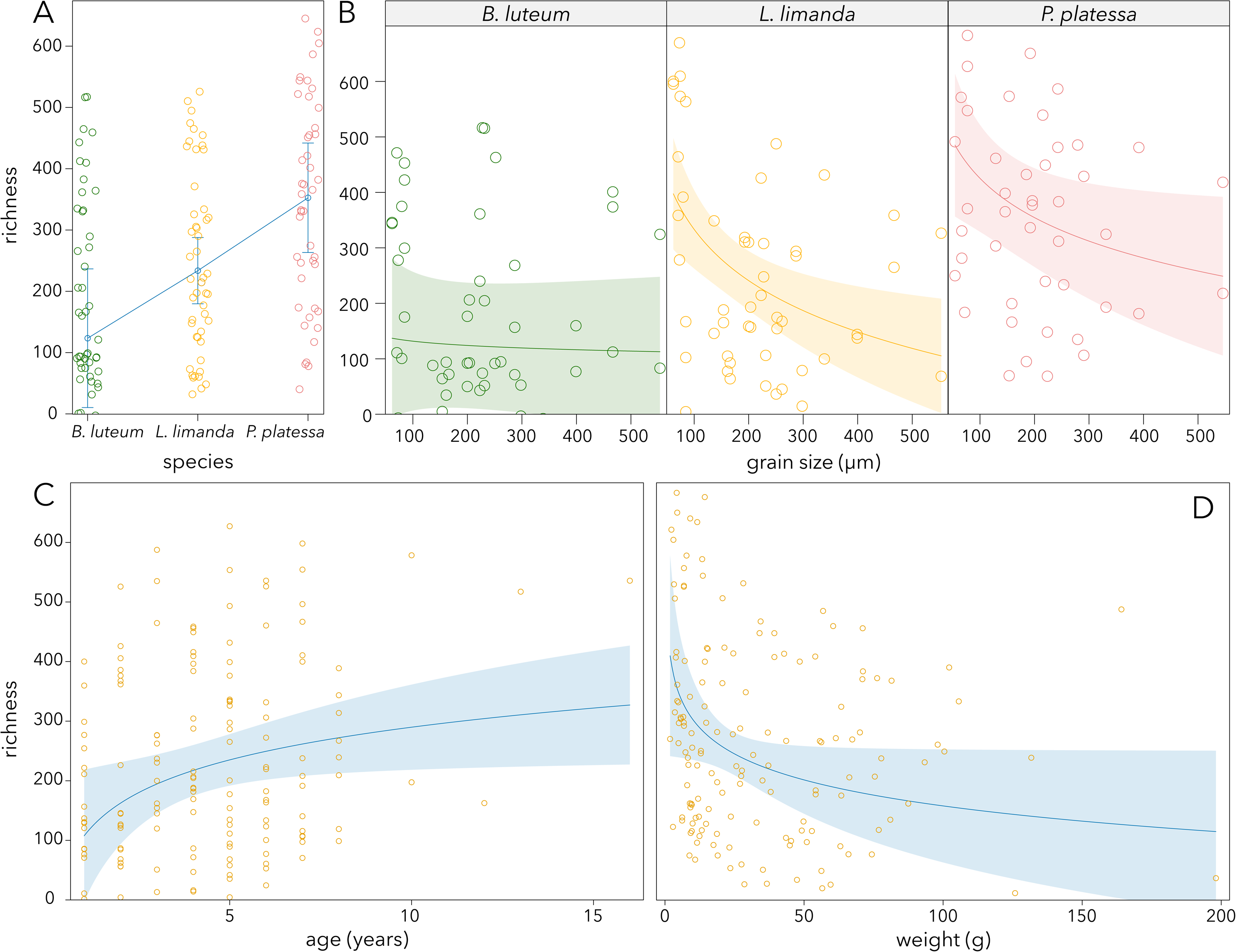
Diversity represented as OTU richness for species, median grain size (separated for species), age and weight.

### 3.2 Community composition

NMDS (based on Bray-Curtis distances) did not reveal strong clustering patterns, but indicated that community composition varied with age, condition factor, median grain size and trawling intensity (Figure4A), and subtly differed among species (Figure 4B).

**Figure 4:**
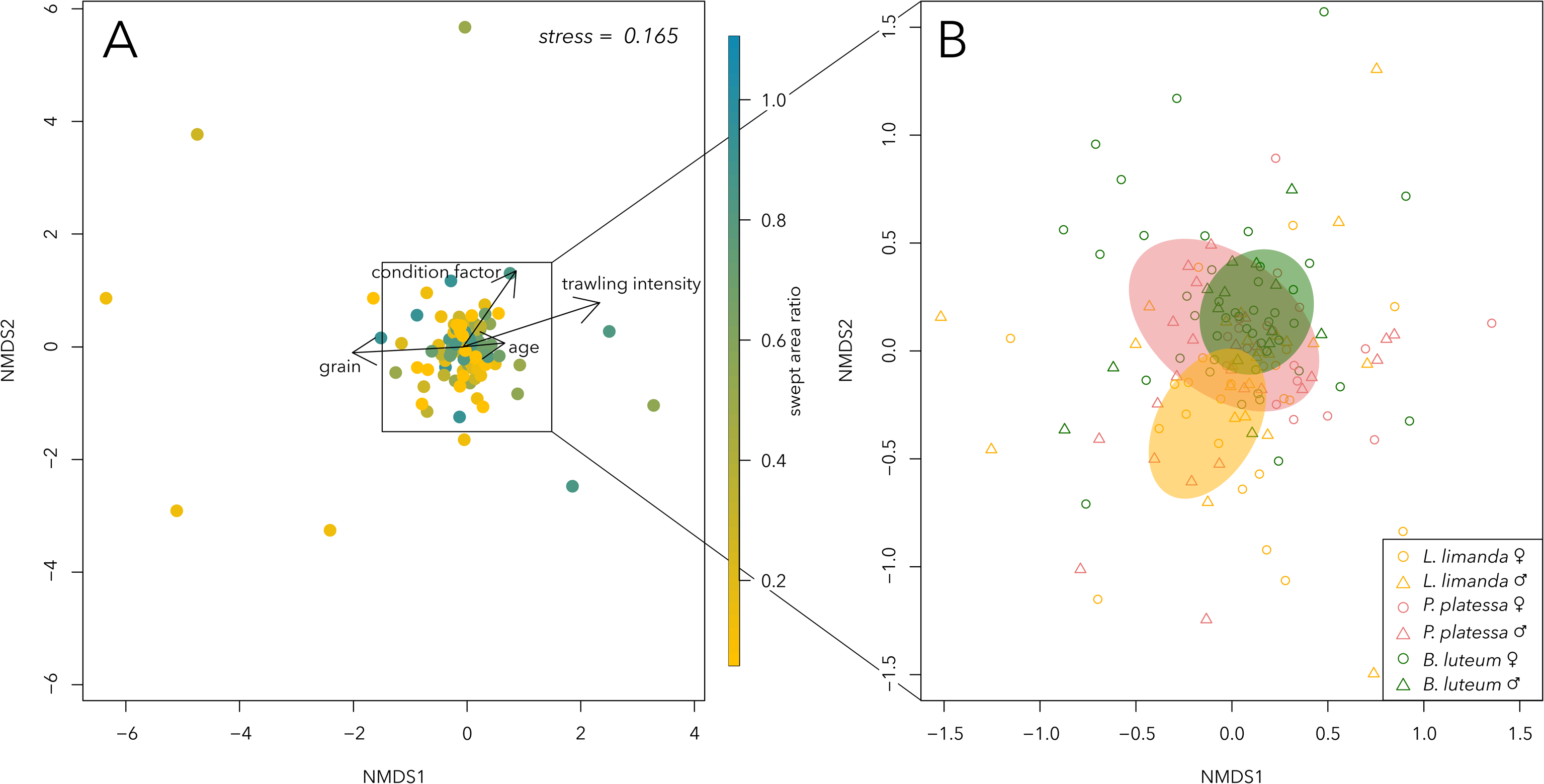
nMDS plots based on Bray-Curtis distances display compositional dissimilarities of the intestinal microbial communities. Variables that significantly explain dissimilarities in community composition are shown for continuous variables and displayed as vectors (A) and for species (B), with ellipses drawn around the centroids based on the standard deviation of the data points.

These patterns were confirmed by the PERMANOVA analysis, which could explain only 16.9% of the overall variation in community composition, but resolved the variables species (F_2,141_ = 1.678, *p*□<□0.001, R^2^ = 0.020), condition factor (F_1,141_ = 1.561, *p*□=□0.015, R^2^ = 0.015), the interaction between condition factor and species (F_2,141_ = 1.311, *p*□=□0.036, R^2^ = 0.036), age (F_2,141_ = 1.834, *p* = 0.003, R^2^ = 0.011), median grain size (F_1,141_ = 3.621, *p*□<□0.001, R^2^ = 0.021) and trawling intensity (F_1,141_ = 3.488, *p*□<□0.001, R^2^ = 0.021) as significant predictors of community composition.

### 3.3 Differential abundances

The differential abundance analysis (Figure 5A) identified various OTUs that were negatively or positively associated with trawling intensity (negative: 29, positive: 24, Figure 5B), and median grain size (negative: 12, positive 35, Figure 5C). The host variable with the highest amount of differentially abundant OTUs was the condition factor (negative: 13, positive: 11, Figure 5D), followed by age (negative: 3, positive: 10, Figure 5E). For *B. luteum* (Figure 5F), *L. limanda* (Figure 5G), and *P. platessa* (Figure 5H), the relative abundance of 6, 10, and 10 OTUs, respectively, increased exclusively with each species. Additionally, 16 OTUs showed significantly lower relative abundance in *B. luteum*, 12 in *L. limanda*, and 2 in *P. platessa*.

**Figure 5:**
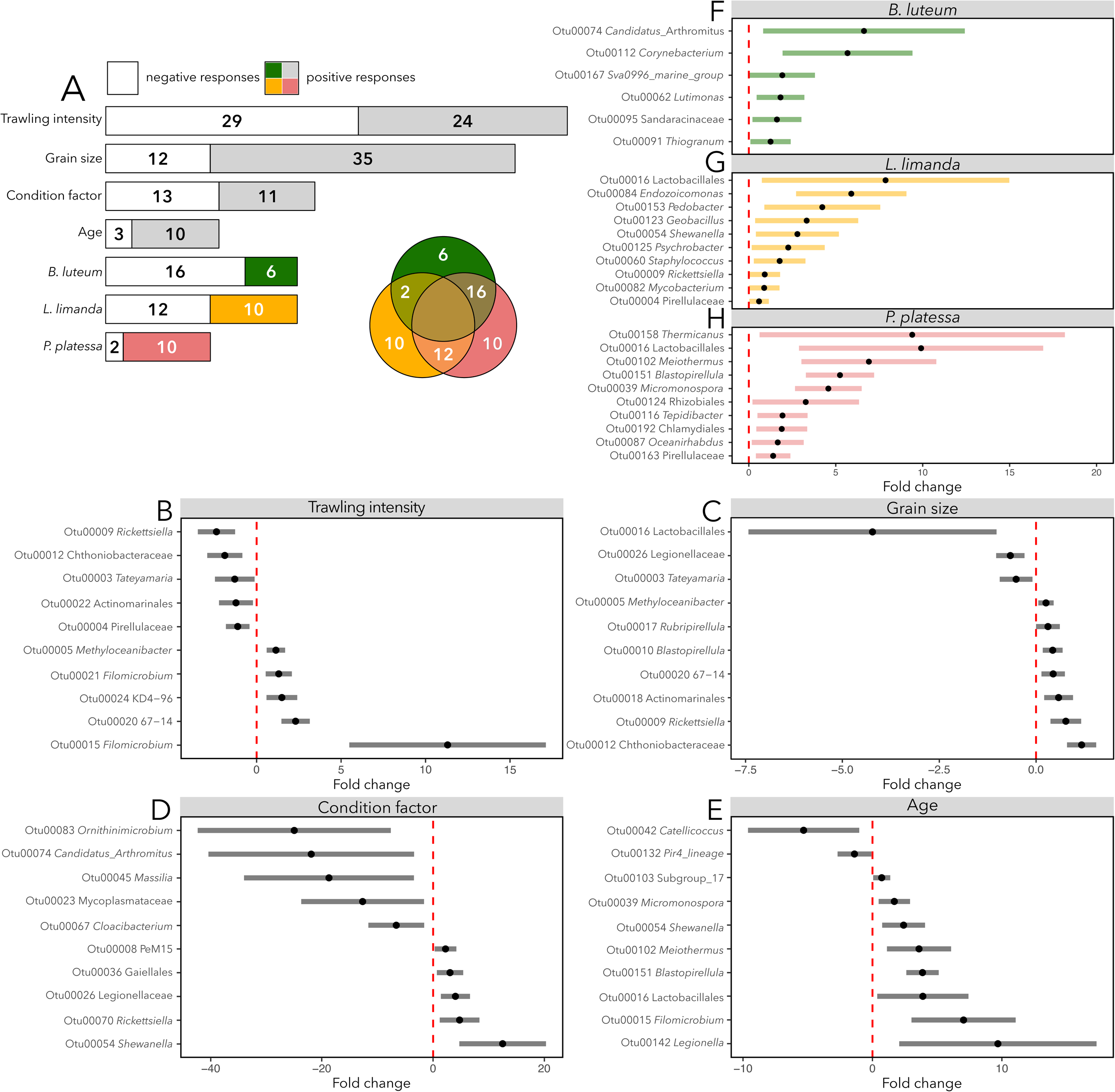
(A) The number of OTUs that significantly differ in abundance, either positively (grey or colored) or negatively (white) in response to environmental and host predictors. Positive responses of differentially abundant OTUs are displayed separately for each species in a Venn diagram, where overlapping areas indicate the number of OTUs that show an increase in abundance in both species. In forest plots, the differentially abundant OTUs in relation to trawling intensity (B), median grain size (C), condition factor (D), age (E) and species (F-H) are illustrated. For each differentially abundant OTU, a negative fold change represents a negative response and a positive fold change value a positive response. The length of the bars indicates the 95% confidence intervals.

## 4. Discussion

This study provides new insights into the intestinal microbial community of three flatfish species from the Southeastern North Sea, highlighting the influence of environmental and host predictors on bacterial diversity and community composition. Overall, the predictors evaluated in this study were able to account for a portion of the total variation, but our models also indicated substantial variation to remain unexplained, indicating that other variables, not examined in the present study, importantly contribute as well to microbial composition and diversity in the fish intestines. The significance of sediment properties (i.e., measured by the median grain size) and trawling intensity emphasizes the important role of environmental factors, which is consistent with previous studies (Hovda *et al*. 2012; Leray *et al*. 2021; MacFarlane *et al*. 1986; Ramírez & Romero 2017; Smith *et al*. 2015). Additionally, we identified several host variables (species, age, condition factor and weight) that contribute to diversity as well as to community composition. However, while we expected that intestinal microbiota would vary between sexes our study found no significant differences between females and males on either diversity or community composition, suggesting that other factors, such as environmental influences or host genetics, play a more dominant role in shaping the microbial community.

It is important to emphasize that an interplay of different factors shapes the intestinal microbial community (Egerton *et al*. 2018; Kanika *et al*. 2025; Xie *et al*. 2024). Some environmental variables such as temperature (Givens 2014; Neuman *et al*. 2016), salinity (Dehler *et al*. 2017; Hieu *et al*. 2022; Lozupone & Knight 2007), water microbiota (Giatsis *et al*. 2015; Xiong *et al*. 2019), pollutants (Adamovsky *et al*. 2018; Mulcahy 2002; Spilsbury *et al*. 2022; Suzzi *et al*. 2022) or diet (Bolnick et al., 2014; Li et al., 2017; Liu et al., 2016a; Xia et al., 2014), were not considered in this study, although they are known to affect the intestinal microbiota of fish. Diet especially, is known to have a primary influence on the diversity and composition of intestinal microbiota (Bolnick et al., 2014; Li et al., 2017; Liu et al., 2016a; Xia et al., 2014). While variation in diet may be partially captured by both host and environmental variables, we did not have direct information of dietary intake of individual flatfishes, which makes other dietary factors likely candidate sources for the variation that this study could not explain.

The presence of several similar bacterial taxa in the intestinal microbiota of one or more fish species from different populations suggests that these taxa play an important role in host intestinal functions (Roeselers *et al*. 2011). As reviewed in the study by Rombout et al. (2011), the phylum Pseudomonadota is the most common in the intestinal microbial community in fish, as it is in *B. luteum*, *L. limanda*, and *P. platessa* in this study comprising about 29% of all reads. Another review by Ghanbari et al. (2015) found that the phyla Pseudomonadota, Bacteroidetes, and Bacillota make up to 90% of the fish intestinal microbiota. For the here studied flatfish species, these phyla account for only ∼40% of all amplicon reads. Instead, the phyla Actinomycetota and Planctomycetota make up a significantly larger share, with the most abundant three phyla accounting for around 70% of the reads found.

### 4.1 Intestinal microbiota vary among flatfish species

It is well known that the intestinal microbiota are species dependent and often vary even among closely related species (Miyake et al. 2015; Navarrete et al. 2012; Smith et al. 2015; Xie et al. 2024). For instance, Miyake et al. (2015) showed that intestinal microbial community composition differed even among closely related species within the same genus (*Acanthurus*). The three species analyzed in this study are more distantly related, belonging to the same order (Pleuronectiformes): *P. platessa* and *L. limanda* are both members of the family Pleuronectidae, while *B. luteum* belongs to the family Soleidae (Ahyong et al. 2023). Given that evolutionary distance is positively correlated with differences in intestinal microbiota, it is expected that these species have distinct intestinal microbiota (Li et al., 2017). Moreover, Smith et al. (2015) examined the intestinal microbiota of different populations of threespine stickleback (*Gasterosteus aculeatus*) and found that its composition and diversity varied, with more genetically divergent populations having more distinct intestinal microbiota. Our results are in line with this, showing that both community composition and OTU-richness varied among the three flatfish species. In addition, we found that changes in composition with the condition factor were species-specific, and changes in OTU richness with sediment type were different among species.

Our study identified 6 OTUs that were significantly more abundant in *B. luteum*, and 10 OTUs in both *P. platessa* and *L. limanda*. Interestingly, one of the *L. limanda* specific OTUs was classified to the genus *Endozoicomonas*, which is recognized as a diverse symbiont, found across a wide range of marine animals, including sponges, corals, mollusks, and fish (Neave et al. 2016). While the flatfish species studied here are benthic predators, there are subtle dietary differences. A study by Schückel et al. (2012), examined the dietary overlap among four flatfish species, including the flatfish studied here, throughout the German Bight and found differences in prey selection. This prey resource partitioning among species likely contributes to the differences in prokaryotic community composition and diversity found here among species. Beyond species-specific dietary differences, other unique traits may further contribute to the distinct composition of the intestinal microbiota. Even though all three species live in marine demersal habitats, only *L. limanda* and *P. platessa* tolerate brackish water. Since salinity is an important factor influencing the composition of the microbiota this may contribute to differences found in intestinal community composition within species (Dehler et al. 2017; Hieu et al. 2022; Lozupone & Knight 2007). Further, there are likely differences in migratory behavior among the flatfishes studied here (Rijnsdorp *et al*. 1992; Marriott *et al*. 2016), as it influences both physiological changes and environmental exposure, and may therefore also contribute to differences among species (Llewellyn *et al*. 2016; Hamilton *et al*. 2019; Liu *et al*. 2021).

### 4.2 Microbiota change with age in composition and become more diverse

Microbial community assembly of the intestine in fish can be divided into colonization and persistence (Smith *et al*. 2015). The interplay between colonization and persistence begins immediately after hatching when microbes start to colonize the intestine (Blanch *et al*. 1997; Lauzon *et al*. 2010). In addition to the skin or gills, the intestine acts as an entry point, allowing prokaryotes in the surrounding environment and the first food to colonize it (Legrand *et al*. 2020). With each successive food or water intake, the intestinal microbiota of the fish becomes more different from the environment (Stephens *et al*. 2016) and diversifies further (Hansen & Olafsen 1999). Accordingly, the diversity of intestinal microbiota tends to increase with age (Li *et al*. 2017; Llewellyn *et al*. 2016; Zhang *et al*. 2018). However, Yan *et al*. (2016) found that diversity decreased with age in freshwater fish species. Especially in early life phases, environmental factors can have a particularly strong and long-lasting impact on the composition of the microbiota (Stephens *et al*. 2016). Considering that fish are exposed to various factors throughout their life that can influence their bacterial community, it is natural to expect a gradual and consistent change with age. Our results are in line with many other studies, who found that microbial diversity in the intestines increases with age in fishes (Ringoe & Birkbeck 1999; Zhang *et al*. 2018), as well as in humans (Nayak, 2010). In the studied flatfish species, we found a similar increasing trend in OTU richness (but not on the effective species number). Furthermore, community composition changed with age, and we identified 6 OTUs that were significantly less abundant while 7 OTUs decreased with age.

### 4.3 Intestinal microbiota do not differ between females and males

Our data showed no evidence for differences in diversity between sexes. These findings are somewhat contrasting to other studies on fish intestinal microbiota where more pronounced differences have been detected between females and males (Chen *et al*. 2022; Liu *et al*. 2016a; Piazzon *et al*. 2019). In a study by Martyniuk et al. (2022) a comparison between females and males in zebrafish found also no differences in diversity between sexes, but several individual taxa were more common in either males or females. For example, male zebrafish showed higher abundance in the families Erythrobacteraceae and Lamiaceae (Martyniuk *et al*. 2022). In another study by Li et al. (2016) on largemouth bronze gudgeon (*Coreius guichenoti*), certain taxa were found to differ in abundance between sexes. Pseudomonadota was the most abundant phylum in males, whereas females exhibited dominance of five different phyla. Additionally, Li et al. (2016) found significant differences in diversity. Such sex-associated differences may be explained by dietary variations between females and males (Bolnick *et al*. 2014), but could also be driven by physiological factors, such as differences in hormone production (Liu *et al*. 2016b; He *et al*. 2021).

### 4.4 Richness decreases with size and composition changes

Few studies have examined the relationship between size and the intestinal microbiota in fish. Since length and weight were highly correlated, we used weight as a proxy for size. Size is linked to other physiological variables such as length or age. Even though a fish’s weight increases with age, especially in early life stages, growth rate reduces with age. This is one of the reasons that makes size and age different from each other and potentially impact the intestinal microbiota differently. Our study was able to disentangle these effects to some extent, as we found that OTU-richness increased with age, whereas a decreasing trend in richness was found for weight, after correcting for age. This result, however, is contrasting with the general observation made across vertebrates (Xie *et al*. 2024) that larger organisms, which consequently have a longer intestine, exhibiting higher intestinal microbial diversity. Likewise, a study by Zhao et al. (2023) on *Gymnocypris chilianensis* found that diversity was higher in larger individuals (>300g) than in smaller ones. These results align with the island biogeography theory (MacArthur & Wilson 2001), which has been linked to diversity of intestinal microbiota, predicting that as an isolated ecosystem, the intestine, can support greater diversity as it increases in size (Ramos Sarmiento *et al*. 2024). Considering the different habitats in the intestines that are associated with distinct microbial communities, this correlation gets even more evident (McCallum & Tropini 2023). First of all, the intestine is composed of the foregut, midgut, and hindgut, each hosting distinct microbial communities (Egerton *et al*. 2018; Hovda *et al*. 2007; Ktari *et al*. 2012; Minich *et al*. 2022; Ringø *et al*. 2006). Furthermore, allochthonous microbiota are transient and associated with digesta, while autochthonous microbiota colonize the mucosal surface, forming the core community Interestingly, our study suggests a size-diversity relationship for flatfish intestinal microbiota that is opposite from what island theory predicts. Exceptions, however, exist as for instance a study on largemouth bronze gudgeon found no effect of weight on diversity (Li et al. 2016).

### 4.5 Fulton’s condition factor is associated with compositional changes

In this study, we demonstrated a significant effect of the variable “condition factor”, as well as an interaction between species and condition factor on community composition. However, no significant effect was found on diversity. Among all studied host variables, condition factor was found to have most differently abundant OTUs consisting of 13 negatively differently abundant OTUs and 11 positively differently abundant OTUs. While weight provides a measure of size, the condition factor, as a ratio of weight to length, offers a relative measure that reflects a fish’s health and food availability (Heino & Kaitala 1999). Also in this case, diet, which is known as an important factor influencing the composition of the intestinal microbiota across vertebrates, and specifically in fish, may be associated with the condition factor. In particular, food availability, or more precisely periods of starvation, are a prime example that affects the condition factor and the intestinal microbiota simultaneously. Consequently, Xia et al. (2014) observed shifts in bacterial communities in Asian seabass, with a significant enrichment of Bacteroidetes and a significant depletion of Betaproteobacteria as a result of starvation. Although comparable studies that analyze the relationship between the condition factor and the intestinal microbiota in fish are rare, community composition of the human intestinal microbiota has been reported to vary with the body mass index, which can be considered equivalent to Fulton’s condition factor (Dominianni *et al*. 2015; Goodrich *et al*. 2014). The relationship between the intestinal microbiota and the condition factor is reciprocal, as some prokaryotes of the intestinal microbiota are able to break down complex sugars and provide essential short-chain fatty acids and energy as well as other nutrients, directly influencing the nutritional and health condition of the fish (Talwar *et al*. 2018). Due to this fundamental importance of the microbiota for the condition of a fish, the intestinal microbial diversity has been used a biomarker for fish health and metabolic capacity (Xiong *et al*. 2019).

### 4.6 Microbial composition and diversity vary with sediment type

With their bottom-dwelling way of life and their food, which largely consists of organisms living in the sediment (Schückel *et al*. 2012), flatfish are in close contact with both the sediment and its specific microbiota. Sediment properties (i.e., median grain size, mud content and organic matter content) are primary drivers of microbial community composition and diversity of the top sediment layer (Bonthond *et al*. 2023). Moreover, also the organisms that make up the flatfish diet and pass through their intestines vary across habitat types and are thus strongly dependent on sediment properties (Huys *et al*. 1992; Neumann *et al*. 2017; Reiss *et al*. 2010). Using the median grain size as a simple measure for sediment type (e.g., mud, sand or gravel), we found that microbial diversity and composition within the flatfish intestine is indeed linked with local sediment properties.

That the sediment is an important source of microbiota found in the intestine is also suggested by others. Although grass carp primarily inhabit the mid to upper water column, Wu et al. (2012) found that their intestinal microbiota composition mainly originates from the surrounding water and sediment. Species dwelling closer to the bottom sediment, such as flatfish, may be even more influenced by it. Besides the highly significant association with OTU-richness and community composition, the median grain size also yielded a large number of differentially abundant OTUs, with 47 OTUs decreasing with median grain size, and 35 OTUs increasing.

### 4.7 Intestinal microbial community composition varies with trawling intensity

We found that trawling intensity explained small but significant changes in intestinal community composition. In total, we detected 24 OTUs to increase with trawling intensity, while 29 OTUs decreased. While trawling activity does not impact the isolated microbial ecosystems in the intestines of individual flatfishes directly, we propose three potential indirect pathways that may explain this observed trend.

First, mobile bottom-contacting fishing gears impact the environment physically. Beam trawls penetrate several centimeters deep into the seafloor, resuspending large amounts of sediment and organic matter, and modifying the seabed morphology (Puig *et al*. 2012). This affects both the water column and the seafloor, with which flatfish live in close contact.

Second, trawling alters benthic faunal communities. In the North Sea, up to 70% of benthic invertebrates, including bivalves, polychaetes, echinoderms or ophiuroids, die when they get dragged by a trawl (reviewed in Eigaard *et al*. 2017, Sciberras *et al*. 2018). Many of these animals naturally influence the remineralization of organic matter and the regeneration of nutrients by microorganisms (Hooper *et al*. 2005; Olsgard *et al*. 2008), but also make up the flatfish diet, which importantly impacts intestinal microbiota composition and diversity (Ringø *et al*. 2006; Xie *et al*. 2024). Link et al. (2002) analyzed dietary data of flatfish in the Northwest Atlantic over 25 years and found that the average weight of stomach contents of flatfish decreased in heavily fished areas. This supports that trawling can influence the diet of flatfishes. Moreover, besides changes in the identity of organisms in the flatfish diet, periods of reduced food availability and starvation are known to alter the intestinal microbial community (Xia *et al*. 2014). Therefore, trawling driven changes in flat fish diet may impact the intestinal microbiota in different ways and present a possible cause of the here observed association between intestinal microbial community composition and trawling intensity.

Third, the demersal flatfish species studied here live in close contact with the sediment, which also acts as a source of microbes that colonize their intestines (Wu *et al*. 2012). Sediment microbiota have been shown to vary with bottom trawling effort as well, showing a decrease in alpha diversity and change in overall community composition (Bruce *et al*. 2022; Bonthond *et al*. 2023). Therefore, trawling related shifts in benthic microbiota may offer another potential pathway through which trawling intensity could indirectly impact the intestinal microbial community of flatfish.

While physical alterations to the seabed morphology, changes in faunal communities and therefore diet, and shifts in sediment microbiota, present interesting hypotheses for how trawling may indirectly affect flat fish intestinal microbiota, they currently remain speculative. Nonetheless, they currently offer the best explanation for the observed association between bottom trawling intensity and intestinal microbiota composition. Moreover, they highlight the need for further investigation and serve as a hypothetical basis for future research on the effects of environmental disturbance caused by fishing activities on fish health and fish intestinal microbiota in particular.

## 5 Conclusions

Here, we disentangled how species identity, age, size, condition factor, and environmental factors (i.e., sediment type and trawling intensity), contribute to shaping the intestinal microbiota of the demersal flatfish species *B. luteum*, *L. limanda* and *P. platessa* in the southeastern North Sea. The strong effects of sediment type and trawling intensity indicate that the environment plays a determinant role in shaping the intestinal microbiota of these flatfishes. To the best of our knowledge, this study is the first to detect an association with bottom trawling intensity. This adds on previous work, that found benthic microbiota to vary along a trawling gradient (Bonthond *et al*. 2023), and may hint that such effects could extent to higher trophic levels. However, we note that substantial variation could not be explained by our models, indicating that other processes, not captured by the variables examined here, importantly influence fish intestinal microbiota as well. Dietary factors likely account for a substantial portion of the unexplained variation in both composition and diversity of the flatfish intestinal microbiota. Another important factor that we were not able to account for is the migratory history of the flatfish studied (Marriott *et al*. 2016; Liu *et al*. 2021).

While many studies have focused on captive fish due to their economic importance in aquaculture, wild populations remain understudied (Kanika *et al*. 2025). Given that Ramírez & Romero (2017) found differences in intestinal microbial communities between wild and captive Fine Flounder (*Paralichthys adspersus*), and Xie et al. (2024) documented such differences across vertebrates in a meta-analysis, the focus of this study on wild populations contributes to narrowing this knowledge gap. Furthermore, this study helps to compensate for a geographic bias, as most studies on this topic to date have come from North America or East Asia, while Central Europe and Africa tend to be underrepresented (Kanika *et al*. 2025).

These findings contribute to our understanding of how host variables as well as environmental and anthropogenic processes may directly or indirectly affect host-associated microbial communities, with potential ecological and evolutionary implications. As this study is the first to observe an association of trawling effort and microbial community composition of flatfish intestines, it merits for more research to identify the mechanisms that underly this trend, gain insight into the long-term impacts, and potential consequences for fish health and population dynamics.

## Acknowledgements

We thank the crew of the RV Solea for all their support during the fieldwork. Birgit Brinkmann and Petra Schwarz are thanked for their help at the ICBM lab. We are particular grateful to Valeria Adrian-Schütte and Jana Bäger (Thünen Institute of Sea Fisheries) for determining the age of individual fishes through otolith analysis. Further, we express our gratitude to Jessica Van der Maesen for illustrating the flatfish species depicted in Figure 1. This is publication number 106 that uses data from the Senckenberg am Meer Metabarcoding and SNG laboratory.

This study was funded by the German Federal Ministry of Education and research (BMBF), through DAM: MGF North Sea I and II (Grant numbers 03F0847B and 03F0936C).

## Data availability

The de-multiplexed V4-16S gene amplicon reads and associated metadata are available from the European Nucleotide Archive under the Bioproject accession number PRJEB88596. Data and R-scripts used for the analyses are available on GitHub at https://github.com/gbonthond/flatfish_microbiota.

## Author contributions

HH, HN, PJS and GB conceptualized the study. Field collections were conducted by HH, HN and GB. MG, SK and GB conducted laboratory work. MG and GB processed data and carried out the formal analysis. MG and GB drafted the manuscript. All authors contributed to revising the manuscript.

## Ethics statement

Animals sampled for this study were obtained from trawling catches collected during regular monitoring by the Thünen Institute of Sea Fisheries aboard the German Research Vessel Solea, which is operated by the Federal Office for Agriculture and Food (BLE), the authority responsible for regulating fishing activities in German waters. Thereby, fishing was conducted with permission from German authorities. Since the animals experienced no additional stress beyond standard commercial fishing practices, no further authorization or ethics approval was required. The species studied are neither protected by legislation nor classified as threatened or endangered. All research complied with the European directive 2010/63/EU on the protection of animals used for scientific purposes.

## References

Adamovsky, O., Buerger, A.N., Wormington, A.M., Ector, N., Griffitt, R.J., Bisesi, J.H., et al. (2018). The gut microbiome and aquatic toxicology: An emerging concept for environmental health: The microbiome and aquatic toxicology. Environ. Toxicol. Chem., 37, 2758–2775.

Ahyong, S., Boyko, C.B., Bailly, N., Bernot, J., Bieler, R., Brandão, S.N., et al. (2023). World Register of Marine Species (WoRMS).

Amoroso, R.O., Pitcher, C.R., Rijnsdorp, A.D., McConnaughey, R.A., Parma, A.M., Suuronen, P., et al. (2018). Bottom trawl fishing footprints on the world’s continental shelves. Proc. Natl. Acad. Sci., 115.

Anderson, M.J. (2001). A new method for non-parametric multivariate analysis of variance: NON-PARAMETRIC MANOVA FOR ECOLOGY. Austral Ecol., 26, 32–46.

Arun, D. & Midhun, S.J. (2023). Microbiome of fish. In: Recent Advances in Aquaculture Microbial Technology. Elsevier, pp. 15–33.

Bates, D., Mächler, M., Bolker, B. & Walker, S. (2015). Fitting Linear Mixed-Effects Models Using lme4. J. Stat. Softw., 67.

Blanch, A.R., Alsina, M., Simón, M. & Jofre, J. (1997). Determination of bacteria associated with reared turbot (**Scophthalmus maximus**) larvae. J. Appl. Microbiol., 82, 729–734.

Bolnick, D.I., Snowberg, L.K., Hirsch, P.E., Lauber, C.L., Org, E., Parks, B., et al. (2014). Individual diet has sex-dependent effects on vertebrate gut microbiota. Nat. Commun., 5, 4500.

Bonthond, G., Beermann, J., Gutow, L., Neumann, A., Rafael Barboza, F., Desiderato, A., et al. (2023). Benthic microbial biogeographic trends in the North Sea are shaped by an interplay of environmental drivers and bottom trawling effort. ISME Commun.

Bruce, S.A., Aytur, S.A., Andam, C.P. & Bucci, J.P. (2022). Metagenomics to characterize sediment microbial biodiversity associated with fishing exposure within the Stellwagen Bank National Marine Sanctuary. Sci. Rep., 12, 9499.

Chen, Z.-W., Jin, X.-K., Gao, F.-X., Gui, J.-F., Zhao, Z. & Shi, Y. (2022). Comparative analyses reveal sex-biased gut microbiota in cultured subadult pufferfish Takifugu obscurus. Aquaculture, 558, 738366.

De La Cuesta-Zuluaga, J., Kelley, S.T., Chen, Y., Escobar, J.S., Mueller, N.T., Ley, R.E., et al. (2019). Age- and Sex-Dependent Patterns of Gut Microbial Diversity in Human Adults. mSystems, 4, e00261–19.

Dehler, C.E., Secombes, C.J. & Martin, S.A.M. (2017). Seawater transfer alters the intestinal microbiota profiles of Atlantic salmon (Salmo salar L.). Sci. Rep., 7, 13877.

Dickey-Collas, M., Nash, R.D.M., Brunel, T., Payne, M.R., Corten, A., Geffen, A.J., et al. (2010). Lessons learned from stock collapse and recovery of North Sea herring: a review. Int. Counc. Explor. Sea.

Dinan, T.G. & Cryan, J.F. (2016). Mood by microbe: towards clinical translation.

Dominianni, C., Sinha, R., Goedert, J.J., Pei, Z., Yang, L., Hayes, R.B., et al. (2015). Sex, Body Mass Index, and Dietary Fiber Intake Influence the Human Gut Microbiome. PLOS ONE, 10, e0124599.

Egerton, S., Culloty, S., Whooley, J., Stanton, C. & Ross, R.P. (2018). The Gut Microbiota of Marine Fish. Front. Microbiol., 9, 873.

Eigaard, O.R., Bastardie, F., Hintzen, N.T., Buhl-Mortensen, L., Buhl-Mortensen, P., Catarino, R., et al. (2017). The footprint of bottom trawling in European waters: distribution, intensity, and seabed integrity. ICES J. Mar. Sci., 74, 847–865.

Fan, G., Song, Y., Yang, L., Huang, X., Zhang, S., Zhang, M., et al. (2020). Initial data release and announcement of the 10,000 Fish Genomes Project (Fish10K). GigaScience, 9.

FAO. (2022). The State of World Fisheries and Aquaculture 2022 - Towards Blue Transformation. FAO - Food and Agriculture Organisation of the United Nations, Rome.

FAO. (2024). The State of World Fisheries and Aquaculture 2024 – Blue Transformation in action. Rome.

Froese, R. & Pauly, Y. (2024). Species in the North Sea.

Ghanbari, M., Kneifel, W. & Domig, K.J. (2015). A new view of the fish gut microbiome: Advances from next-generation sequencing. Aquaculture, 448, 464–475.

Ghotbi, M., Kelting, O., Blümel, M. & Tasdemir, D. (2022). Gut and Gill-Associated Microbiota of the Flatfish European Plaice (Pleuronectes platessa): Diversity, Metabolome and Bioactivity against Human and Aquaculture Pathogens. Mar. Drugs, 20, 573.

Giatsis, C., Sipkema, D., Smidt, H., Heilig, H., Benvenuti, G., Verreth, J., et al. (2015). The impact of rearing environment on the development of gut microbiota in tilapia larvae. Sci. Rep., 5, 18206.

Givens, C.E. (2014). A Fish Tale: Comparison of the Gut Microbiome of 15 Fish Species and the Influence of Diet and Temperature on its Composition. Dep. Mar. Sci. Univ. Ga. USA, 232.

Gohl, D.M., Vangay, P., Garbe, J., MacLean, A., Hauge, A., Becker, A., et al. (2016). Systematic improvement of amplicon marker gene methods for increased accuracy in microbiome studies. Nat. Biotechnol., 34, 942–949.

Goodrich, J.K., Waters, J.L., Poole, A.C., Sutter, J.L., Koren, O., Blekhman, R., et al. (2014). Human Genetics Shape the Gut Microbiome. Cell, 159, 789–799.

Hamilton, E.F., Element, G., Van Coeverden De Groot, P., Engel, K., Neufeld, J.D., Shah, V., et al. (2019). Anadromous Arctic Char Microbiomes: Bioprospecting in the High Arctic. Front. Bioeng. Biotechnol., 7, 32.

Hansen, G.H. & Olafsen, J.A. (1999). Bacterial Interactions in Early Life Stages of Marine Cold Water Fish. Microb. Ecol., 38, 1–26.

He, S., Li, H., Yu, Z., Zhang, F., Liang, S., Liu, H., et al. (2021). The Gut Microbiome and Sex Hormone-Related Diseases. Front. Microbiol., 12, 711137.

Heino & Kaitala. (1999). Evolution of resource allocation between growth and reproduction in animals with indeterminate growth. J. Evol. Biol., 12, 423–429.

Hieu, D.Q., Hang, B.T.B., Lokesh, J., Garigliany, M.-M., Huong, D.T.T., Yen, D.T., et al. (2022). Salinity significantly affects intestinal microbiota and gene expression in striped catfish juveniles. Appl. Microbiol. Biotechnol., 106, 3245– 3264.

Holmlund, C.M. & Hammer, M. (1999). Ecosystem services generated by fish populations. Ecol. Econ., 29, 253–268.

Hooper, D.U., Chapin, F.S., Ewel, J.J., Hector, A., Inchausti, P., Lavorel, S., et al. (2005). EFFECTS OF BIODIVERSITY ON ECOSYSTEM FUNCTIONING: A CONSENSUS OF CURRENT KNOWLEDGE. Ecol. Monogr., 75, 3–35.

Hovda, M.B., Fontanillas, R., McGurk, C., Obach, A. & Rosnes, J.T. (2012). Seasonal variations in the intestinal microbiota of farmed Atlantic salmon (Salmo salar L.): Seasonal variations in the intestinal microbiota of Salmo salar L. Aquac. Res., 43, 154–159.

Hovda, M.B., Lunestad, B.T., Fontanillas, R. & Rosnes, J.T. (2007). Molecular characterisation of the intestinal microbiota of farmed Atlantic salmon (Salmo salar L.). Aquaculture, 272, 581–588.

Huang, Q., Sham, R.C., Deng, Y., Mao, Y., Wang, C., Zhang, T., et al. (2020). Diversity of gut microbiomes in marine fishes is shaped by host-related factors. Mol. Ecol., 29, 5019–5034.

Huys, R., Herman, P.M.J., Heip, C.H.R. & Soetaert, K. (1992). The meiobenthos of the North Sea: density, biomass trends and distribution of copepod communities. ICES J. Mar. Sci., 49, 23–44.

ICES. (2022). Greater North Sea ecoregion – fisheries overview, 13045887 Bytes.

Jost, L. (2006). Entropy and diversity. Oikos, 113, 363–375.

Kaiser, M.J., Collie, J.S., Hall, S.J., Jennings, S. & Poiner, I.R. (2002). Modification of marine habitats by trawling activities: prognosis and solutions. F H F H E R E S.

Kanika, N.H., Liaqat, N., Chen, H., Ke, J., Lu, G., Wang, J., et al. (2025). Fish gut microbiome and its application in aquaculture and biological conservation. Front. Microbiol., 15, 1521048.

Klindworth, A., Pruesse, E., Schweer, T., Peplies, J., Quast, C., Horn, M., et al. (2013). Evaluation of general 16S ribosomal RNA gene PCR primers for classical and next-generation sequencing-based diversity studies. Nucleic Acids Res., 41, e1–e1.

Ktari, N., Jridi, M., Bkhairia, I., Sayari, N., Ben Salah, R. & Nasri, M. (2012). Functionalities and antioxidant properties of protein hydrolysates from muscle of zebra blenny (Salaria basilisca) obtained with different crude protease extracts. Food Res. Int., 49, 747–756.

Lauzon, H.L., Gudmundsdottir, S., Petursdottir, S.K., Reynisson, E., Steinarsson, A., Oddgeirsson, M., et al. (2010). Microbiota of Atlantic cod (Gadus morhua L.) rearing systems at pre- and posthatch stages and the effect of different treatments: Microbiota of cod rearing systems. J. Appl. Microbiol., no-no.

Legrand, T.P.R.A., Wynne, J.W., Weyrich, L.S. & Oxley, A.P.A. (2020). A microbial sea of possibilities: current knowledge and prospects for an improved understanding of the fish microbiome. Rev. Aquac., 12, 1101–1134.

Leray, M., Wilkins, L.G.E., Apprill, A., Bik, H.M., Clever, F., Connolly, S.R., et al. (2021). Natural experiments and long-term monitoring are critical to understand and predict marine host–microbe ecology and evolution. PLOS Biol., 19, e3001322.

Li, T., Long, M., Li, H., Gatesoupe, F.-J., Zhang, X., Zhang, Q., et al. (2017). Multi-Omics Analysis Reveals a Correlation between the Host Phylogeny, Gut Microbiota and Metabolite Profiles in Cyprinid Fishes. Front. Microbiol., 8.

Li, X., Yan, Q., Ringø, E., Wu, X., He, Y. & Yang, D. (2016). The influence of weight and gender on intestinal bacterial community of wild largemouth bronze gudgeon (Coreius guichenoti, 1874). BMC Microbiol., 16, 191.

Link, J.S., Bolles, K. & Milliken, C.G. (2002). The Feeding Ecology of Flatfish in the Northwest Atlantic. J. Northwest Atl. Fish. Sci., 30, 1–17.

Liu, H., Guo, X., Gooneratne, R., Lai, R., Zeng, C., Zhan, F., et al. (2016a). The gut microbiome and degradation enzyme activity of wild freshwater fishes influenced by their trophic levels. Sci. Rep., 6, 24340.

Liu, Y., Li, X., Li, J. & Chen, W. (2021). The gut microbiome composition and degradation enzymes activity of black Amur bream (*Megalobrama terminalis*) in response to breeding migratory behavior. Ecol. Evol., 11, 5150–5163.

Liu, Y., Yao, Y., Li, H., Qiao, F., Wu, J., Du, Z., et al. (2016b). Influence of Endogenous and Exogenous Estrogenic Endocrine on Intestinal Microbiota in Zebrafish. PLOS ONE, 11, e0163895.

Llewellyn, M.S., McGinnity, P., Dionne, M., Letourneau, J., Thonier, F., Carvalho, G.R., et al. (2016). The biogeography of the atlantic salmon (*Salmo salar*) gut microbiome. ISME J., 10, 1280–1284.

Lozupone, C.A. & Knight, R. (2007). Global patterns in bacterial diversity. Proc. Natl. Acad. Sci., 104, 11436–11440.

MacArthur, R.H. & Wilson, E.O. (2001). The theory of island biogeography. Princeton landmarks in biology. 13th printing and first Princeton landmarks in biology ed. Princeton university press, Princeton Oxford.

MacFarlane, R.D., McLaughlin, J.J. & Bullock, G.L. (1986). Quantitative and Qualitative Analysis of Gut Flora in Striped Bass from Estuarine and Coastal Marine Habitats. J. Wildl. Dis., 22, 344–348.

Marriott, A., McCarthy, I., Ramsay, A. & Chenery, S. (2016). Discriminating nursery grounds of juvenile plaice (Pleuronectes platessa) in the south-eastern Irish Sea using otolith microchemistry. Mar. Ecol. Prog. Ser., 546, 183–195.

Martyniuk, C.J., Buerger, A.N., Vespalcova, H., Rudzanova, B., Sohag, S.R., Hanlon, A.T., et al. (2022). Sex-dependent host-microbiome dynamics in zebrafish: Implications for toxicology and gastrointestinal physiology. Comp. Biochem. Physiol. Part D Genomics Proteomics, 42, 100993.

McCallum, G. & Tropini, C. (2023). The gut microbiota and its biogeography. Nat. Rev. Microbiol.

Minich, J.J., Härer, A., Vechinski, J., Frable, B.W., Skelton, Z.R., Kunselman, E., et al. (2022). Host biology, ecology and the environment influence microbial biomass and diversity in 101 marine fish species. Nat. Commun., 13, 6978.

Miyake, S., Ngugi, D.K. & Stingl, U. (2015). Diet strongly influences the gut microbiota of surgeonfishes. Mol. Ecol., 24, 656–672.

Mulcahy, M.F. (2002). Diseases of flatfish. Department of Zoology and Animal Ecology, University College Cork, Ireland.

Nash, R.D.M., Valencia, A.H. & Geffen, A.J. (2006). The origin of Fulton’s condition factor - Setting the record straight.

Navarrete, P., Magne, F., Araneda, C., Fuentes, P., Barros, L., Opazo, R., et al. (2012). PCR-TTGE Analysis of 16S rRNA from Rainbow Trout (Oncorhynchus mykiss) Gut Microbiota Reveals Host-Specific Communities of Active Bacteria. PLoS ONE, 7, e31335.

Nayak, S.K. (2010). Role of gastrointestinal microbiota in fish: Role of gastrointestinal microbiota in fish. Aquac. Res., 41, 1553–1573.

Neave, M.J., Apprill, A., Ferrier-Pagès, C. & Voolstra, C.R. (2016). Diversity and function of prevalent symbiotic marine bacteria in the genus Endozoicomonas. Appl. Microbiol. Biotechnol., 100, 8315–8324.

Neuman, C., Hatje, E., Zarkasi, K.Z., Smullen, R., Bowman, J.P. & Katouli, M. (2016). The effect of diet and environmental temperature on the faecal microbiota of farmed Tasmanian Atlantic Salmon (*Salmo salar* L.). Aquac. Res., 47, 660– 672.

Neumann, H., Diekmann, R., Emeis, K.-C., Kleeberg, U., Moll, A. & Kröncke, I. (2017). Full-coverage spatial distribution of epibenthic communities in the south-eastern North Sea in relation to habitat characteristics and fishing effort. Mar. Environ. Res., 130, 1–11.

Nie, L., Zhou, Q.-J., Qiao, Y. & Chen, J. (2017). Interplay between the gut microbiota and immune responses of ayu (Plecoglossus altivelis) during Vibrio anguillarum infection. Fish Shellfish Immunol., 68, 479–487.

NOAH. (2015). NOAH Habitat atlas portal: Porosity of Marine Sediments.

Oksanen, J., Simpson, G.L., Blanchet, F.G., Kindt, R., Legendre, P., Minchin, P.R., et al. (2013). vegan: Community Ecology Package.

Olsgard, F., Schaanning, M.T., Widdicombe, S., Kendall, M.A. & Austen, M.C. (2008). Effects of bottom trawling on ecosystem functioning. J. Exp. Mar. Biol. Ecol., 366, 123–133.

OSPAR. (2017). OSPAR Bottom Fishing Intensity - Surface & Subsurface.

Piazzon, M.C., Naya-Català, F., Simó-Mirabet, P., Picard-Sánchez, A., Roig, F.J., Calduch-Giner, J.A., et al. (2019). Sex, Age, and Bacteria: How the Intestinal Microbiota Is Modulated in a Protandrous Hermaphrodite Fish. Front. Microbiol., 10, 2512.

Puig, P., Canals, M., Company, J.B., Martín, J., Amblas, D., Lastras, G., et al. (2012). Ploughing the deep sea floor. Nature, 489, 286–289.

Quast, C., Pruesse, E., Yilmaz, P., Gerken, J., Schweer, T., Yarza, P., et al. (2012). The SILVA ribosomal RNA gene database project: improved data processing and web-based tools. Nucleic Acids Res., 41, D590–D596.

Ramírez, C. & Romero, J. (2017). Fine Flounder (Paralichthys adspersus) Microbiome Showed Important Differences between Wild and Reared Specimens. Front. Microbiol., 08.

Ramos Sarmiento, K., Carr, A., Diener, C., Locey, K.J. & Gibbons, S.M. (2024). Island biogeography theory provides a plausible explanation for why larger vertebrates and taller humans have more diverse gut microbiomes. ISME J., 18, wrae114.

Ray, A.K., Ringoe, E. & Ghosh, K. (2012). Enzymeproducing bacteria isolated from fish gut: a review. Aquac. Nutr., 18, 465–492.

Reiss, H., Degraer, S., Duineveld, G.C.A., Kröncke, I., Aldridge, J., Craeymeersch, J.A., et al. (2010). Spatial patterns of infauna, epifauna, and demersal fish communities in the North Sea. ICES J. Mar. Sci., 67, 278–293.

Rijnsdorp, A.D., Vethaak, A.D. & van Leeuwen, P.I. (1992). Population biology of dab Limanda limanda in the southeastern North Sea. Mar. Ecol. Prog. Ser., 91, 19–35.

Ringø, E., Sperstad, S., Myklebust, R., Refstie, S. & Krogdahl, Å. (2006). Characterisation of the microbiota associated with intestine of Atlantic cod (Gadus morhua L.). Aquaculture, 261, 829–841.

Ringoe, E. & Birkbeck, T. (1999). Intestinal microflora of fish larvae and fry. Aquac. Res., 30, 73–93.

Roeselers, G., Mittge, E.K., Stephens, W.Z., Parichy, D.M., Cavanaugh, C.M., Guillemin, K., et al. (2011). Evidence for a core gut microbiota in the zebrafish. ISME J., 5, 1595–1608.

Rolig, A.S., Mittge, E.K., Ganz, J., Troll, J.V., Melancon, E., Wiles, T.J., et al. (2017). The enteric nervous system promotes intestinal health by constraining microbiota composition. PLOS Biol., 15, e2000689.

Rombout, J.H.W.M., Abelli, L., Picchietti, S., Scapigliati, G. & Kiron, V. (2011). Teleost intestinal immunology (online first). Fish Shellfish Immunol. 2010.

Saba, G.K., Burd, A.B., Dunne, J.P., Hernández-León, S., Martin, A.H., Rose, K.A., et al. (2021). Toward a better understanding of fish-based contribution to ocean carbon flux. Limnol. Oceanogr., 66, 1639–1664.

Schloss, P.D., Westcott, S.L., Ryabin, T., Hall, J.R., Hartmann, M., Hollister, E.B., et al. (2009). Introducing mothur: Open-Source, Platform-Independent, Community-Supported Software for Describing and Comparing Microbial Communities. Appl. Environ. Microbiol., 75, 7537–7541.

Schückel, S., Sell, A.F., Kröncke, I. & Reiss, H. (2012). Diet overlap among flatfish species in the southern North Sea. J. Fish Biol., 80, 2571–2594.

Sciberras, M., Hiddink, J.G., Jennings, S., Szostek, C.L., Hughes, K.M., Kneafsey, B., et al. (2018). Response of benthic fauna to experimental bottom fishing: A global meta-analysis. Fish Fish., 19, 698–715.

Smith, C.C.R., Snowberg, L.K., Gregory Caporaso, J., Knight, R. & Bolnick, D.I. (2015). Dietary input of microbes and host genetic variation shape among-population differences in stickleback gut microbiota. ISME J., 9, 2515–2526.

Spilsbury, F., Foysal, M.J., Tay, A. & Gagnon, M.M. (2022). Gut Microbiome as a Potential Biomarker in Fish: Dietary Exposure to Petroleum Hydrocarbons and Metals, Metabolic Functions and Cytokine Expression in Juvenile Lates calcarifer. Front. Microbiol., 13, 827371.

Stephens, W.Z., Burns, A.R., Stagaman, K., Wong, S., Rawls, J.F., Guillemin, K., et al. (2016). The composition of the zebrafish intestinal microbial community varies across development. ISME J., 10, 644–654.

Suzzi, A.L., Stat, M., MacFarlane, G.R., Seymour, J.R., Williams, N.LR., Gaston, T.F., et al. (2022). Legacy metal contamination is reflected in the fish gut microbiome in an urbanised estuary. Environ. Pollut., 314, 120222.

Talwar, C., Nagar, S., Lal, R. & Negi, R.K. (2018). Fish Gut Microbiome: Current Approaches and Future Perspectives. Indian J. Microbiol., 58, 397–414.

Tarnecki, A.M., Burgos, F.A., Ray, C.L. & Arias, C.R. (2017). Fish intestinal microbiome: diversity and symbiosis unravelled by metagenomics. J. Appl. Microbiol., 123, 2–17.

Viana, D.F., Zamborain-Mason, J., Gaines, S.D., Schmidhuber, J. & Golden, C.D. (2023). Nutrient supply from marine small-scale fisheries. Sci. Rep., 13, 11357.

Wang, Y., Naumann, U., Wright, S. & Warton, D. (2012). Wang Y, Naumann U, Wright ST, Warton DI.. mvabund - an R package for model-based analysis of multivariate abundance data. Methods Ecol Evol 3: 471–474. *Methods Ecol. Evol.*, 3, 471.

Westcott, S.L. & Schloss, P.D. (2017). OptiClust, an Improved Method for Assigning Amplicon-Based Sequence Data to Operational Taxonomic Units. mSphere, 2, e00073–17.

Wilkins, L.G.E., Leray, M., O’Dea, A., Yuen, B., Peixoto, R.S., Pereira, T.J., et al. (2019). Host-associated microbiomes drive structure and function of marine ecosystems. PLOS Biol., 17, e3000533.

Wu, S., Wang, G., Angert, E.R., Wang, W., Li, W. & Zou, H. (2012). Composition, Diversity, and Origin of the Bacterial Community in Grass Carp Intestine. PLoS ONE, 7, e30440.

Xia, J.H., Lin, G., Fu, G.H., Wan, Z.Y., Lee, M., Wang, L., et al. (2014). The intestinal microbiome of fish under starvation.

Xie, Y., Xu, S., Xi, Y., Li, Z., Zuo, E., Xing, K., et al. (2024). Global meta-analysis reveals the drivers of gut microbiome variation across vertebrates. iMetaOmics, 1, e35.

Xiong, J.-B., Nie, L. & Chen, J. (2019). Current understanding on the roles of gut microbiota in fish disease and immunity. Zool. Res., 40, 70–76.

Yan, Q., Li, J., Yu, Y., Wang, J., He, Z., Van Nostrand, J.D., et al. (2016). Environmental filtering decreases with fish development for the assembly of gut microbiota. Environ. Microbiol., 18, 4739–4754.

Zhang, Z., Li, D., Refaey, M.M., Xu, W., Tang, R. & Li, L. (2018). Host Age Affects the Development of Southern Catfish Gut Bacterial Community Divergent From That in the Food and Rearing Water. Front. Microbiol., 9, 495.

Zhao, Z., Zhao, H., Zhang, L., Huang, Z., Ke, H., Liu, Y., et al. (2023). Integrated analysis of how gender and body weight affect the intestinal microbial diversity of Gymnocypris chilianensis. Sci. Rep., 13, 8811.

Zhu, L., Wang, J. & Bahrndorff, S. (2021). Editorial: The Wildlife Gut Microbiome and Its Implication for Conservation Biology. Front. Microbiol., 12, 697499.

