## Supplemental Files for "Flatfish intestinal microbiota depend on various host traits, and vary with sediment type and bottom trawling effort"

**Table S1.** Model selection tables for Effective number of OTUs and richness.

| | condition | log(age) | log(weight) | log2(grain) | sex | species | SAR | species : condition | species : log(age) | species : log(weight) | species : log2(grain) | species : sex | species : SAR | Df | Log Likeliood | AICc | Delta AICc | AICc weight |
|---|---|---|---|---|---|---|---|---|---|---|---|---|---|---|---|---|---|---|
| ENO | **-** | **-** | **-** | **-** | **-** | **-** | **+** | **-** | **-** | **-** | **-** | **-** | **-** | **4** | **-712.5** | **1433.3** | **0.00** | **0.087** |
| | - | - | - | - | - | - | - | - | - | - | - | - | - | 3 | -714.0 | 1434.1 | 0.84 | 0.057 |
| | - | - | - | + | - | - | + | - | - | - | - | - | - | 5 | -712.2 | 1434.7 | 1.45 | 0.042 |
| | - | + | - | - | - | - | + | - | - | - | - | - | - | 5 | -712.3 | 1435.0 | 1.72 | 0.037 |
| | - | - | + | - | - | - | + | - | - | - | - | - | - | 5 | -712.3 | 1435.0 | 1.72 | 0.037 |
| | + | - | - | - | - | - | + | - | - | - | - | - | - | 5 | -712.4 | 1435.2 | 1.92 | 0.033 |
| richness | **-** | **+** | **+** | **+** | **-** | **+** | **-** | **-** | **-** | **-** | **+** | **-** | **-** | **10** | **-1058.1** | **2137.6** | **0.00** | **0.020** |
| | + | + | + | + | - | + | - | + | - | - | + | - | - | 13 | -1054.7 | 2137.9 | 0.27 | 0.017 |
| | - | + | + | + | - | + | + | - | - | - | + | - | - | 11 | -1057.1 | 2138.1 | 0.44 | 0.016 |
| | + | - | - | + | + | + | - | + | - | - | + | + | - | 14 | -1053.6 | 2138.1 | 0.48 | 0.015 |
| | - | - | - | + | - | - | - | - | - | - | - | - | - | 4 | -1064.9 | 2138.1 | 0.51 | 0.015 |
| | - | - | - | + | + | + | - | - | - | - | + | + | - | 11 | -1057.2 | 2138.2 | 0.59 | 0.015 |
| | + | - | - | + | - | + | - | + | - | - | + | - | - | 11 | -1057.3 | 2138.3 | 0.71 | 0.014 |
| | + | + | - | + | - | + | - | + | - | - | + | - | - | 12 | -1056.2 | 2138.5 | 0.87 | 0.013 |
| | - | - | - | + | + | - | - | - | - | - | - | - | - | 5 | -1064.1 | 2138.5 | 0.89 | 0.013 |
| | + | + | + | + | - | + | + | + | - | - | + | - | - | 14 | -1053.9 | 2138.6 | 1.00 | 0.012 |
| | - | - | - | + | + | + | - | - | - | - | - | + | - | 9 | -1059.7 | 2138.6 | 1.02 | 0.012 |
| | - | - | - | + | - | + | - | - | - | - | + | - | - | 8 | -1060.9 | 2138.7 | 1.12 | 0.011 |
| | + | + | - | + | + | + | - | + | - | - | + | + | - | 15 | -1052.8 | 2138.8 | 1.22 | 0.011 |
| | - | - | - | + | - | - | + | - | - | - | - | - | - | 5 | -1064.3 | 2139.0 | 1.38 | 0.010 |
| | - | - | - | + | - | + | - | - | - | - | - | - | - | 6 | -1063.3 | 2139.1 | 1.44 | 0.010 |
| | - | + | - | + | - | + | - | - | - | - | + | - | - | 9 | -1060.0 | 2139.1 | 1.48 | 0.009 |
| | - | + | - | + | + | + | - | - | - | - | + | + | - | 12 | -1056.6 | 2139.2 | 1.60 | 0.009 |
| | + | + | + | + | + | + | - | + | - | - | + | + | - | 16 | -1051.7 | 2139.2 | 1.61 | 0.009 |
| | - | - | - | + | + | + | + | - | - | - | + | + | - | 12 | -1056.6 | 2139.2 | 1.63 | 0.009 |
| | - | - | - | + | + | - | + | - | - | - | - | - | - | 6 | -1063.4 | 2139.3 | 1.68 | 0.008 |
| | + | - | - | + | - | + | + | + | - | - | + | - | - | 12 | -1056.7 | 2139.4 | 1.83 | 0.008 |
| | - | - | - | + | + | + | + | - | - | - | - | + | - | 10 | -1059.0 | 2139.5 | 1.91 | 0.008 |
| | + | - | - | + | + | + | + | + | - | - | + | + | - | 15 | -1053.1 | 2139.5 | 1.92 | 0.008 |
| | + | + | - | + | - | + | + | + | - | - | + | - | - | 13 | -1055.5 | 2139.5 | 1.94 | 0.007 |

*Abbreviations: Effective Numbers of OTUs (ENO), condition factor (condition), median grain size (grain), swept area ratio (SAR), Akaike information criterion corrected for small sample sizes (AICc), degrees of freedom (Df).*
*Note: Only models with a delta AICc < 2 are shown. Station identity was included as random intercept in all models included in the selection procedure.*

**Table S2.** ANOVA table ENO and richness

| Response | predictor | $\chi^2$ | Df | $Pr(>\chi^2)$[1] | |
|---|---|---|---|---|---|
| ENO | SAR | 0.3055 | 1 | 0.08046 | . |
| Richness | species | 7.3252 | 2 | **0.02567** | * |
| | log(age) | 5.0468 | 1 | **0.02467** | * |
| | log(weight) | 3.8905 | 1 | **0.04855** | * |
| | log2(grain) | 7.7733 | 1 | **0.00530** | ** |
| | species : log2(grain) | 6.7423 | 2 | **0.03435** | * |

*Note: linear mixed effect models with station identity as random intercept*
*[1] Significance codes: p < 0.1 (.), p < 0.05 (*), p < 0.01 (**), p < 0.001 (***)*

**Table S3.** Post-hoc pairwise comparisons on OTU richness

| | Contrast | df | t-value | Pr(>t)[1] | |
|---|---|---|---|---|---|
| species | *B. luteum – L. limanda* | 168 | -1.517 | 0.1311 | |
| | *B. luteum – P. platessa* | 167 | -2.525 | **0.0302** | * |
| | *L. limanda – P. platessa* | 158 | -2.605 | **0.0302** | * |
| species : log2(grain) | *B. luteum* | 96.2 | 2 | 0.7767 | |
| | *L. limanda* | 116.9 | 1 | **0.0052** | ** |
| | *P. platessa* | 137.8 | 2 | 0.0662 | . |

*Note: P values are adjusted with the Holm-method.*
[1] *Significance codes: p < 0.1 (.), p < 0.05 (*), p < 0.01 (**), p < 0.001 (***)*

**Table S4.** PERMANOVA table

| | Df | SumOfSqs | R2 | F | Pr(>F)[1] | |
|---|---|---|---|---|---|---|
| species | 2 | 0.9556 | 0.0198 | 1.6778 | **0.0007** | *** |
| sex | 1 | 0.2175 | 0.0045 | 0.7637 | 0.9088 | |
| condition | 1 | 0.4444 | 0.0092 | 1.5605 | **0.0152** | * |
| log(age) | 1 | 0.5223 | 0.0108 | 1.8341 | **0.0025** | ** |
| log(weight) | 1 | 0.3190 | 0.0066 | 1.1200 | 0.2378 | |
| log2(grain) | 1 | 1.0312 | 0.0213 | 3.6208 | **0.0001** | *** |
| SAR | 1 | 0.9932 | 0.0206 | 3.4877 | **0.0001** | *** |
| species : sex | 2 | 0.6123 | 0.0127 | 1.0758 | 0.2757 | |
| species : condition | 2 | 0.7467 | 0.0155 | 1.3109 | **0.0358** | * |
| species : log(age) | 2 | 0.6433 | 0.0133 | 1.1294 | 0.1740 | |
| species : log(weight) | 2 | 0.5334 | 0.0110 | 0.9366 | 0.6326 | |
| species : log2(grain) | 2 | 0.5645 | 0.0117 | 0.9910 | 0.4788 | |
| species : SAR | 2 | 0.5773 | 0.0119 | 1.0136 | 0.4229 | |
| Residual | 141 | 40.1547 | 0.8311 | | | |
| Total | 161 | 48.3158 | 1.0000 | | | |

*Note: Permanova based on Bray-Curtis distances*
[1] *Significance codes: p < 0.1 (.), p < 0.05 (*), p < 0.01 (**), p < 0.001 (***)*

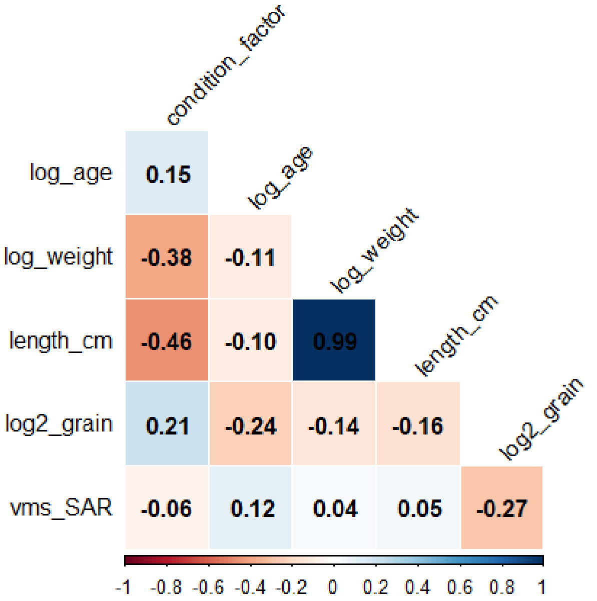


**Figure S1.** Raw spearman rank correlation values among all variables considered in the current study.

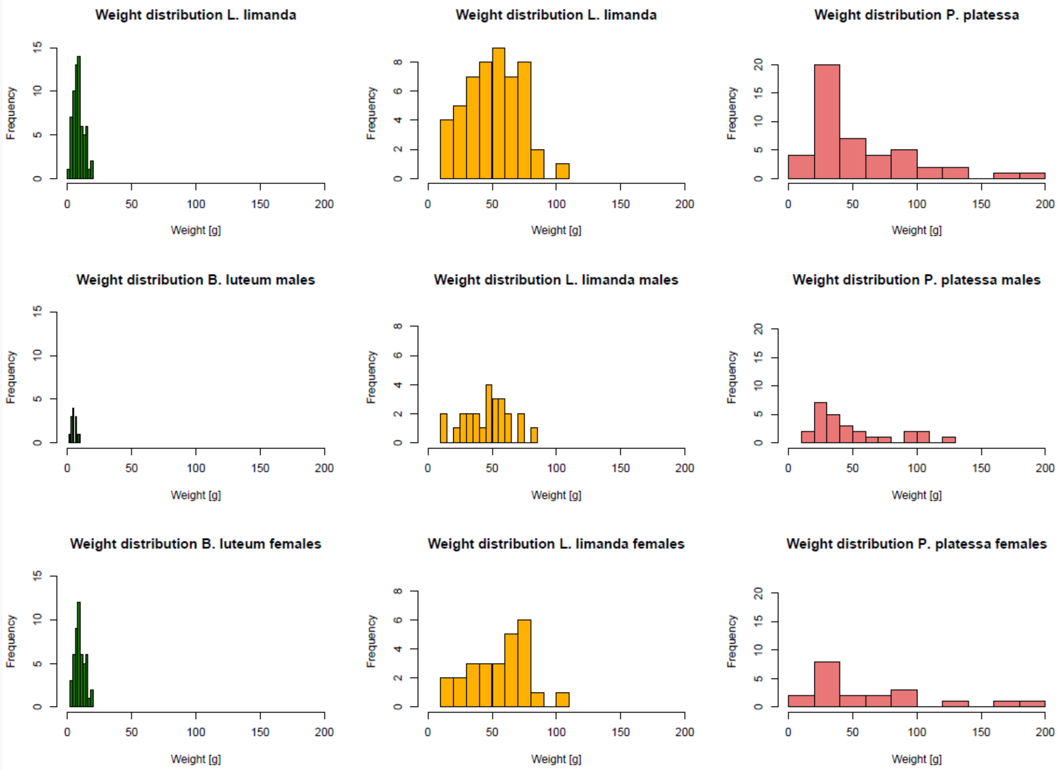


**Figure S2.** Histograms showing weight distribution ranges by species and sex

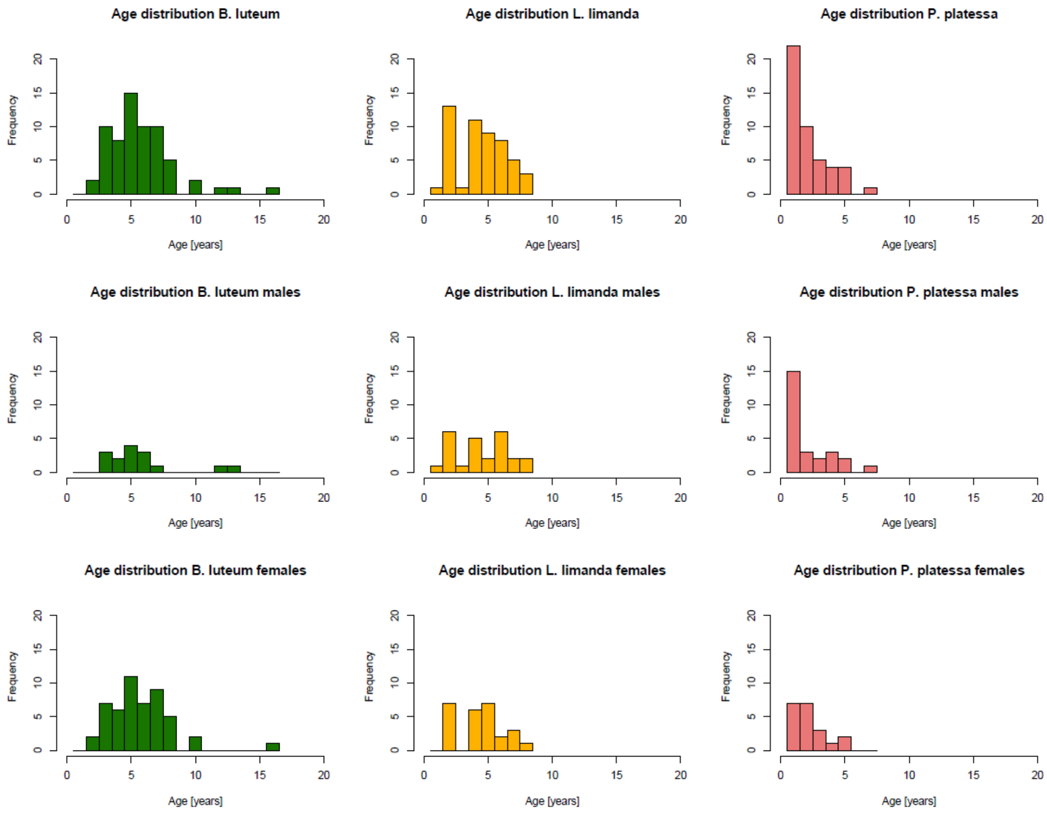


**Figure S3.** Histograms showing age distribution ranges by species and sex

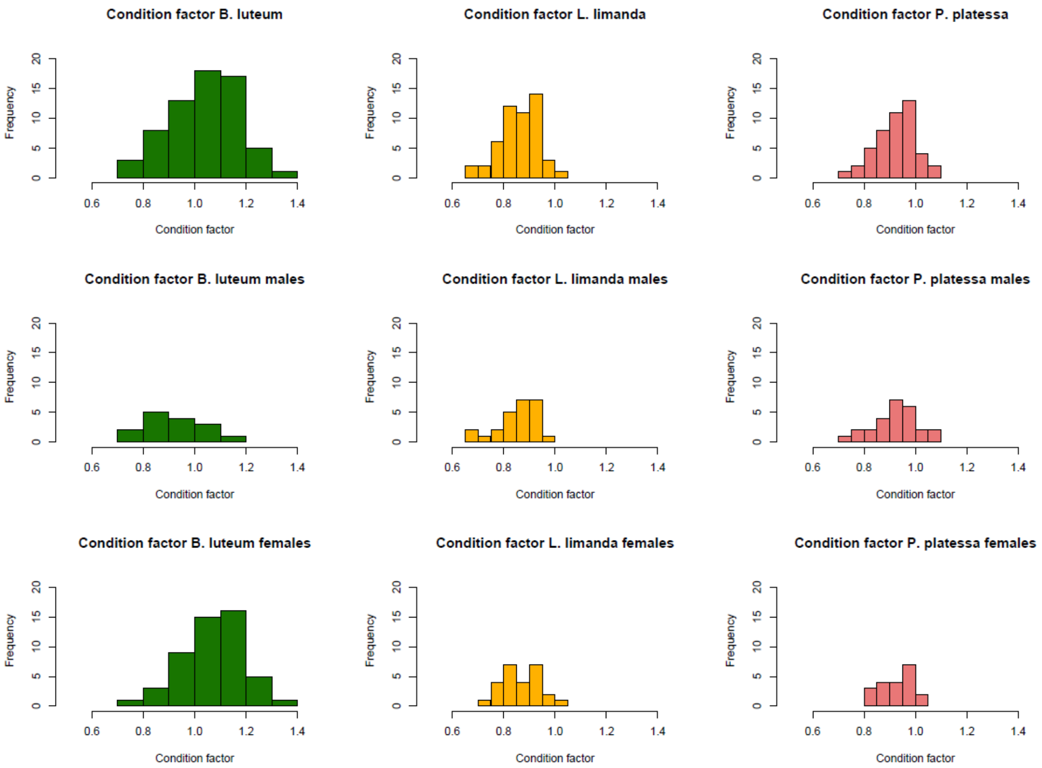


**Figure S4.** Histograms showing condition factor distribution ranges by species and sex

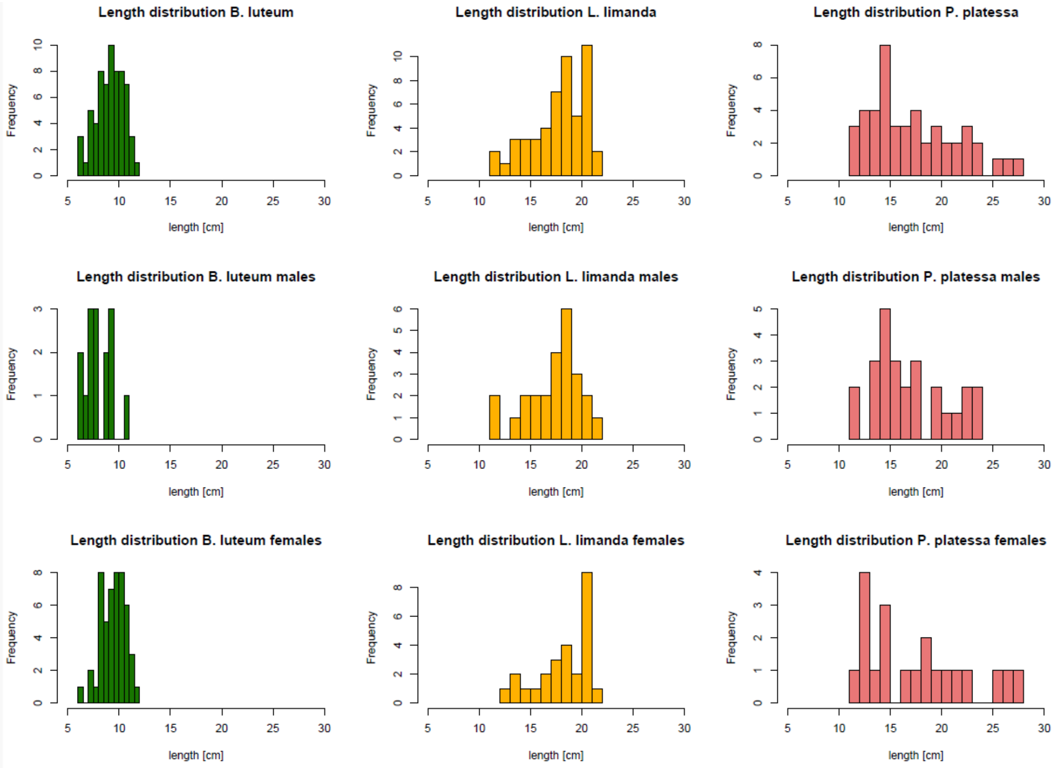


**Figure S5.** Histograms showing length distribution ranges by species and sex
